# PFAS triggers a SpoT-associated metabolic switch that promotes persister-like phenotype in *Salmonella* Typhi

**DOI:** 10.64898/2026.05.12.724561

**Authors:** Debayan Ganguli, Shramana Chakraborty, Shreya Dasgupta, Sameer Ranjan Sahoo, Debjani Bhattacharya, Sudip Dey, Ananda Pal, Ratan Gachhui, Santasabuj Das

## Abstract

Per- and polyfluoroalkyl substances (PFAS) are new pollutants in the environment whose effects on bacteria’s physiology is not well understood. In this study, we show that exposure to PFAS causes membrane depolarization in *Salmonella* enterica serovar Typhi. This works as a metabolic uncoupler that breaks down proton motive force without immediately killing the cells. This disturbance results in a significant elevation of intracellular NADH and NAD⁺ levels while preserving redox equilibrium, signifying an augmented metabolic flux. At the same time, we see that β-oxidation pathways are turned on, which suggests that the cells are shifting toward breaking down fats to make up for the lack of energy.

Even though there are more reducing equivalents, ATP levels go down, which is what happens when respiration is uncoupled. This puts the cells in a state of “pseudo-starvation.” This metabolic stress triggers the SpoT-dependent stringent response, leading to the accumulation of (p)ppGpp. Genetic analysis employing Δ*relA* and Δ*relA*Δ*spoT* mutants confirm that SpoT is necessary for this adaptive response. Functionally, PFAS-treated populations show an enhanced proportion of persister-like cells, which connects exposure to environmental pollutant in the environment to antibiotic tolerance.

Our findings reveal a previously unidentified mechanism by which PFAS alters bacterial metabolism and stress responses, facilitating persistence through membrane depolarization, metabolic reconfiguration, and stringent response activation. This study underscores the potential influence of environmental pollutants on bacterial survival mechanisms and antibiotic resistance.

## Introduction

*Salmonella enterica* serovar Typhi (*S*. Typhi) is a human restricted pathogen that causes typhoid fever, which continues to be a major public health problem in many parts of the world (1, 2). The pathogenicity of the bacteria depends on the coordinated expression of virulence genes encoded by *Salmonella* pathogenicity islands (SPIs), especially SPI-1 and SPI-2 and their secreted effectors transported to the host cells by type three secretion systems. These processes are tightly regulated and very sensitive to signals from the environment and the infected host, which allow the pathogen to switch between different physiological states during infection (3, 4).

*S*. Typhi adapts to stress by generating persisters or becoming biofilm-associated. These changes help the bacteria to become antibiotic resistant and live longer. Bacterial persisters are subpopulations of cells that display phenotypic antibiotic tolerance without heritable resistance (5). A key regulator of this process is the stringent response, mediated by small alarmones, (p)ppGpp or guanosine penta- and tetra-phosphates. In Gram negative bacteria, two enzymes, RelA (hydrolase) and SpoT (bifunctional hydrolase/synthetase) regulate (p)ppGpp synthesis (6). RelA primarily detects amino acid starvation through mechanisms linked to ribosomes, while SpoT responds to a wider range of stress signals, such as fatty acids and carbon source limitation, membrane changes etc. (7, 8). Accumulation of (p)ppGpp leads to global transcriptional reprogramming, often involving induction of the general stress sigma factor RpoS, metabolic downshift, and increased tolerance to antibiotics (9, 10).

Membrane homeostasis and lipid metabolism are progressively acknowledged as essential factors influencing bacterial stress adaptation. Changes in the composition of membranes or the availability of fatty acids can cause compensatory responses, including activation of the pathways for fatty acid biosynthesis and SpoT-dependent signaling (10–12). For example, it was shown that acyl carrier protein and SpoT interact directly to link fatty acid status to (p)ppGpp synthesis (8). These connections imply that membrane-disruptors might serve as significant inducers of persistence via stringent response activation.

Per- and polyfluoroalkyl substances (PFAS) are a group of pollutants, called forever chemicals because of their long persistence in the environment, such as water, soil, and food systems (13). Because of their amphipathic nature and chemical stability, PFAS are known to bind lipid bilayers and change the properties of membranes (14). Although their toxicological effects in eukaryotic systems have been thoroughly investigated, the influence on bacterial physiology, especially in pathogenic organisms, is inadequately understood. New evidence shows that pollutants and other environmental stressors that do not kill bacteria can help them to stick around and form biofilms (15, 16). Nonetheless, a significant knowledge deficiency persists - about whether environmentally pertinent membrane-active pollutants, such as PFAS can directly reconfigure bacterial regulatory networks to induce persistence. More specifically, the degree to which PFAS-induced membrane disruptions activate stringent response signalling, and its impact on the persistence phenotypes remains unexplored.

In this study, we have investigated the impact of PFAS exposure on the physiology of *S*. Typhi, focusing on persistence and biofilm formation. We demonstrate that PFAS elicits a pseudo-starvation response, as indicated by the upregulation of fatty acid catabolism genes, including *acyl coA dehydrogensase*, *fadA* and *fadB* due to low intracellular ATP. This triggers a SpoT-dependent stringent response, resulting in elevated RpoS expression, metabolic reprogramming, and augmented persister cell formation. Our findings designate PFAS as an environmental signal that influences critical regulatory pathways in *S*. Typhi, while at the same time induces membrane depolarization that elicits SpoT-dependent stress adaptation.

## Material and Methods

### Cells and reagents

*Salmonella* Typhi Ty2 strain (ATCC no 19430) and THP-1 cell line (ATCC no TIB 202) were procured from the American Type Culture Collection (ATCC). Cell culture reagents (RPMI, Dulbecco’s modified Eagle’s medium [DMEM], and fetal bovine serum [FBS])) were procured from Invitrogen. Bacterial culture media (Luria-Bertani [LB] broth and LB agar) were purchased from HiMedia. All antibiotics used were procured from Sigma. Ty2Δ*relA* and Ty2Δ*relA* Δ*spoT* mutants were constructed as mentioned in ref 17. PFOA (Cat No 171468), PFNA (Cat No 394459) and PFUnDA (Cat No 446777) were purchased from Sigma.

### PFAS treatment

Ty2 was grown from fresh glycerol stock at 37°C overnight. Next day, the culture was diluted 1:100 with fresh LB media in the presence or absence of PFAS mix (final concentration:10 µM PFOA, 10 µM PFNA and 10 µM PFUnDa) and the culture was grown at 37℃ under shaking condition for 12 hrs. Bacteria recovered by centrifugation was used for all downstream experiments.

### Killing and Re-killing assay

PFAS treated or untreated cultures were diluted in 1:50 ratio (to a final CFU of∼10^8^ CFU/ ml) in fresh LB containing 10 µg/ml Ceftriaxone or 200 µg/ml Azithromycin and incubated at 37℃ under shaking condition with or without PFAS. Cultures were harvested at predetermined time intervals, and the bacteria recovered were washed with 1x PBS, followed by re-suspension in fresh LB. After culturing for 1 hr at 37℃, the bacteria were centrifuged, resuspended in LB and spread on Streptomycin (50 µg/ml) containing LA plates. Bacterial killing was analysed by CFU counts. For re-killing assay, single colony from the plates were grown as above and CFU counts on LA plates were enumerated as described before.

### Biofilm formation

To develop biofilm in test tubes, PFAS treated or untreated Ty2 were diluted in 1:50 ratio (to a final CFU of ∼10^8^ CFU/ ml) in fresh LB, supplemented with 5% Ox-bile. The cultures were incubated at 37℃ for 5 days with or without PFAS. Then, the cultures were discarded and the glass tubes were washed with PBS for five times. The biofilm was stained with crystal violet following the standard protocol. For measuring absorbance measurement, the biofilm was dislodged using ethanol and OD was taken at 570 nm using a plate reader.

To develop biofilm under nutrient deprivation, PFAS treated or untreated Ty2 were diluted in 1:50 ratio (∼10^8^ CFU/ ml) in fresh LB media, diluted with PBS (1:10). The cultures were incubated with or without PFAS in 24 well plates at 37℃ for 5 days under static condition. After discarding the culture, the wells were washed with PBS for five times. The biofilm was stained with crystal violet, followed by absorbance measurement as described above.

To grow biofilm on lettuce leaves, leaves were dipped in 10 % bleach for 3 mins, 100 % ethanol for 3 mins and then washed with autoclaved deionized H_2_O for 15 mins with 3 changes. The leaves were placed into 24 well cell plates and Ty2 biofilms were developed under nutrient starvation as described above. The biofilms grown on the leaves were dislodged using PBS, containing proteinase K and DNase I, followed by vigorous vortexing for 10 mins. The leaves were discarded, bacteria pelleted and plated on Streptomycin (50 µg/ml) containing LA plates at different dilutions.

### EPS quantification

Bacterial cultures, treated with PFAS as mentioned above were diluted at 1:50 with fresh LB and incubated at 37℃ for 5 days in the presence or absence of PFAS. The cultures were centrifuged, and the supernatant was treated with 10 volumes of absolute ethanol at -20℃ overnight. The solution was centrifuged at 12000 rpm for 10 mins. After air-drying the pellets, they were dissolved in 200 µl of water. The resulting EPS was quantified by phenol/ sulphuric acid method. Briefly, 50 µl of the purified EPS was treated with 10 µl phenol and subsequently 500 µl of concentrated sulphuric acid was added. The absorbance of the resulting yellow coloured solution was measured at wavelength of 490 nm.

### Measurement of ATP and NADH

∼10^5^ CFU/ ml PFAS treated or untreated Ty2 were washed with and resuspended in PBS. ATP was measured using BAC Titer-Glo (Promega Cat No G8230) following the manufacturer’s instructions. NADH+NAD^+^ were measured using colorimetric assay kit (Elabscience, Cat No. E-BC-K804-M) following the manufacturer’s instructions. Both ATP and NADH+NAD^+^ were normalised against total CFU.

### Measurement of membrane depolarisation

The fluorescent dye DiBAC4(3) (bis-(1,3-dibutylbarbituric acid) trimethine oxonol) (Merck Cat No D8189) was used to measure the membrane potential. ∼10^5^ CFU/ ml bacterial cells were collected by centrifugation and washed two times with PBS. The cells were resuspended in PBS and stained with DiBAC4(3) at a final concentration of 1µM for 30 minutes at room temperature in the dark. Fluorescence was measured by Flow cytometry (Cytek Aurora) with unstained and untreated cells being included as controls for baseline fluorescence and gating.

### EtBr retention assay

This assay was performed to study efflux pump functions. ∼10^5^ CFU/ ml bacterial cells were incubated with EtBr (1µg/mL) until a stable level of fluorescence was reached. Flow cytometry was used to measure intracellular fluorescence. To evaluate the energy dependence of efflux activity, assays were conducted under two conditions: (i) in the absence of glucose (energy-depleted condition) and (ii) with glucose supplementation (0.2–1%) to replenish cellular energy levels and proton motive force. EtBr retention was calculated as: % EtBr retention = (fluorescence intensity at time t / fluorescence intensity at time 0) × 100.

### qRT PCR

Total RNA was extracted from *Salmonella* enterica serovar Typhi cultures using TRIzol according to the manufacturer’s instructions. RNA samples were treated with DNase I to remove genomic DNA contamination, and RNA integrity was verified by agarose gel electrophoresis. RNA concentration was measured using a spectrophotometer (NanoDrop). First-strand cDNA was synthesized from 1 µg of total RNA using a reverse transcription kit with random hexamers or gene-specific primers following standard protocols. Quantitative PCR was performed using SYBR Green master mix in a real-time PCR system. Each 20 µL reaction contained diluted cDNA, gene-specific primers (0.5 µM each), and SYBR Green reagent. Cycling conditions included initial denaturation at 95°C for 2–3 min, followed by 40 cycles of 95°C for 10–15 s and 60°C for 30 s. Melting curve analysis was performed to confirm amplification specificity. All reactions were performed with at least three biological replicates and three technical replicates, including no-template controls. Gene expression was normalized to a housekeeping gene (gyrA) and relative expression levels were calculated using the ΔΔCt method. Fold change was determined as 2^(−ΔΔCt)^, where ΔCt = Ct(target) − Ct(reference) and ΔΔCt = ΔCt(treated or mutant) − ΔCt(control). Data is presented as mean ± standard deviation (SD), and statistical significance was evaluated using unpaired two-tailed Student’s t-test with p < 0.005 considered significant.

### SMG plate assay

The sensitivity to serine, methionine, and glycine (SMG) was tested by using M9 minimal agar plates enriched with glucose and amino acids. In brief, M9 agar plates with 0.4% glucose, 2 mM MgSO₄, 0.1 mM CaCl₂, and 1.5% agar were prepared with filter sterilized L-serine (100 µg/mL), L-methionine (100 µg/mL), and glycine (100 µg/mL) when the temperature was lowered to around 55°C. The overnight cultures of bacteria were washed twice in sterile 1× PBS to ensure that all the bacteria were depleted from any residual rich medium and then normalised according to cell densities. Serial dilutions of bacteria were plated onto SMG plates and incubated at 37°C for 24–48 h. Growth was observed as colonies next day.

### Statistical analysis

Data are shown as mean ± standard deviation (SD) from at least three biological replicates. Statistical significance was determined using a Mann Whitney Test. A p-value of less than 0.05 was deemed statistically significant.

## Results

### PFAS enhances *Salmonella* Typhi antibiotic persisters

Exposure of *S*. Typhi Ty2 to PFAS (10 µM PFOA, 10 µM PFNA and 10 µM PFUnDa), followed by antibiotics (Ceftriaxone 10µg/ml or Azithromycin 200µg/ml) led to biphasic kinetics of the time-kill curve with significant increase in persister cell formation (Fig. 1 A, D). That increased survival was due to bacterial persisters was confirmed by repeat treatment of the surviving bacteria with antibiotics, showing similar biphasic kinetics, suggesting a transient and heritable persister phenotype (Fig. 1 B, E). Bacterial survival was augmented by 5.2 and 5.6 times after PFAS pre-treatment following exposure to ceftriaxone and azithromycin, respectively for 24h (Fig. 1 C, F), while PFAS exposure alone failed to influence bacterial growth (Fig. S1A). Absence of changes in antibiotic MICs further supported bacterial persister formation (Fig. S1B). Together these results suggest that although PFAS alone does not exert any effect on *S*. Typhi persisters, it significantly enhances antibiotic persister generation.

**Figure 1.**
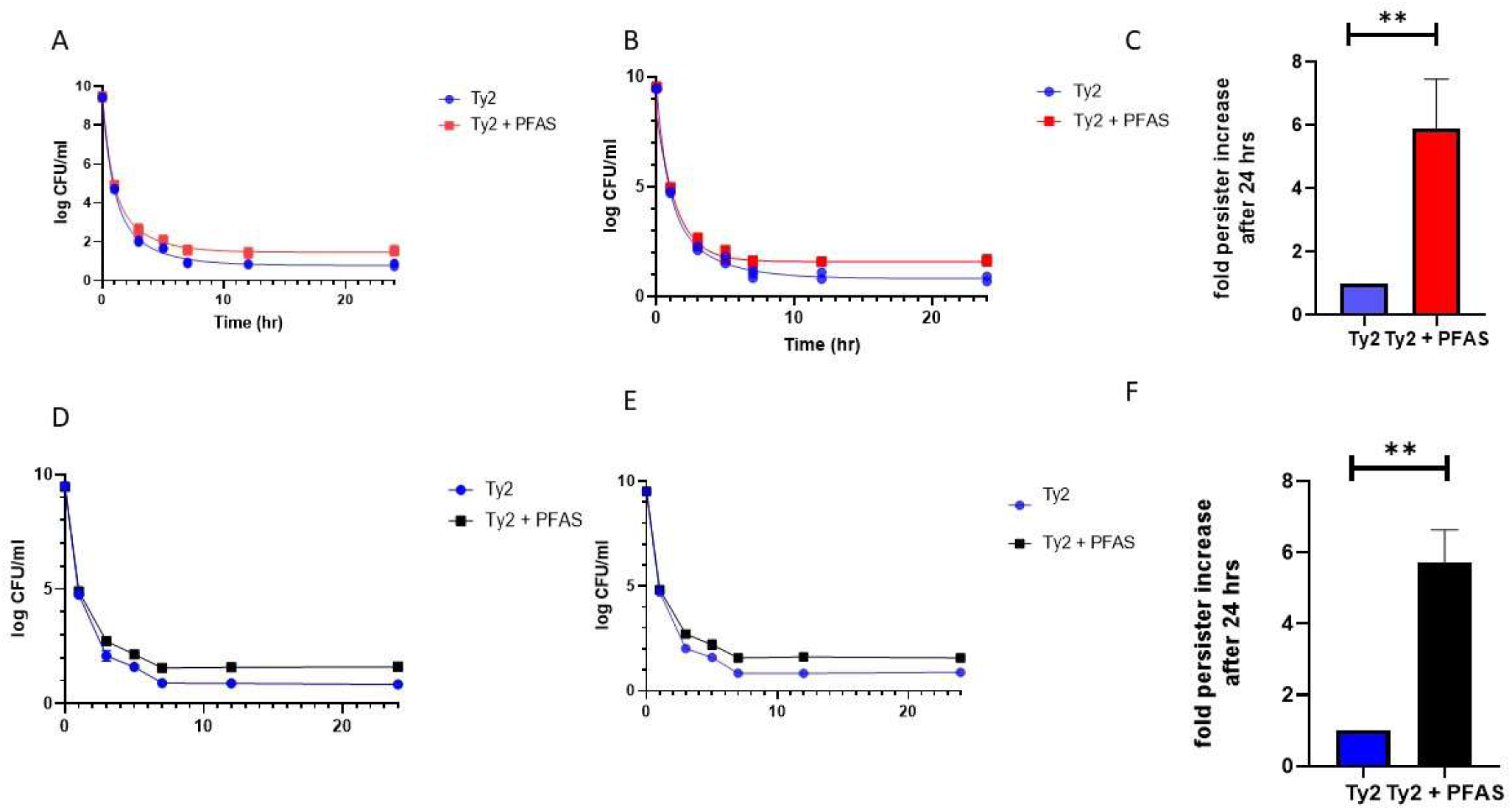
PFAS exposure increases persister population of Ty2 in presence of ceftriaxone and azithromycin. (A) Around 5×10^9^ Ty2 cells were treated or not treated with PFAS and exposed to ceftriaxone (10 µg/mL) in LB at 37°C with shaking. Samples were collected at the time points 1hr, 2hrs, 3hrs, 4 hrs, 5 hrs, 7 hrs and 24 hrs, washed with PBS, resuspended in fresh LB and allowed to recover for 1 h before being plated on LA supplemented with streptomycin (50 µg/mL) for CFU determination. (B) To proof the colonies obtained in (A) are persisters, rekilling assay was performed by culturing individual colonies that had recovered after ceftriaxone treatment overnight and re-exposing them to ceftriaxone under the same conditions, and CFUs were determined as described above. (C) Fold change in persister levels after 24h ceftriaxone exposure calculated from panel (A). (D) Ty2 cells treated or non-treated with PFAS were challenged with azithromycin (100 µg/mL) in a similar manner (A) and surviving populations were quantified over time using the same recovery and plating procedure. (E) Colonies that survived azithromycin exposure were regrown overnight and treated again (100 µg/mL azithromycin) to test persistence stability, then CFUs were quantified. (F) Fold change in persister levels after 24 h treatment with azithromycin (from panel (D). Data represent mean ± SD from three independent biological replicates (n = 3). Statistical significance was determined using a Mann Whitney Test, with *\*\** representing P value < 0.005.

### PFAS increases Salmonella Typhi biofilm formation

Given that bacterial persisters play a critical role in biofilm formation, we checked if PFAS exposure increased *Salmonella* Typhi biofilm. In the presence of nutrients (LB medium, containing 5% ox bile) as well as nutrient deprivation simulated by 1:10 dilution of the LB medium, a significant increase in *S*. Typhi biofilm biomass production was observed after PFAS treatment (Fig. 2 A, B; Fig. S2 A, 2B).

**Figure 2:**
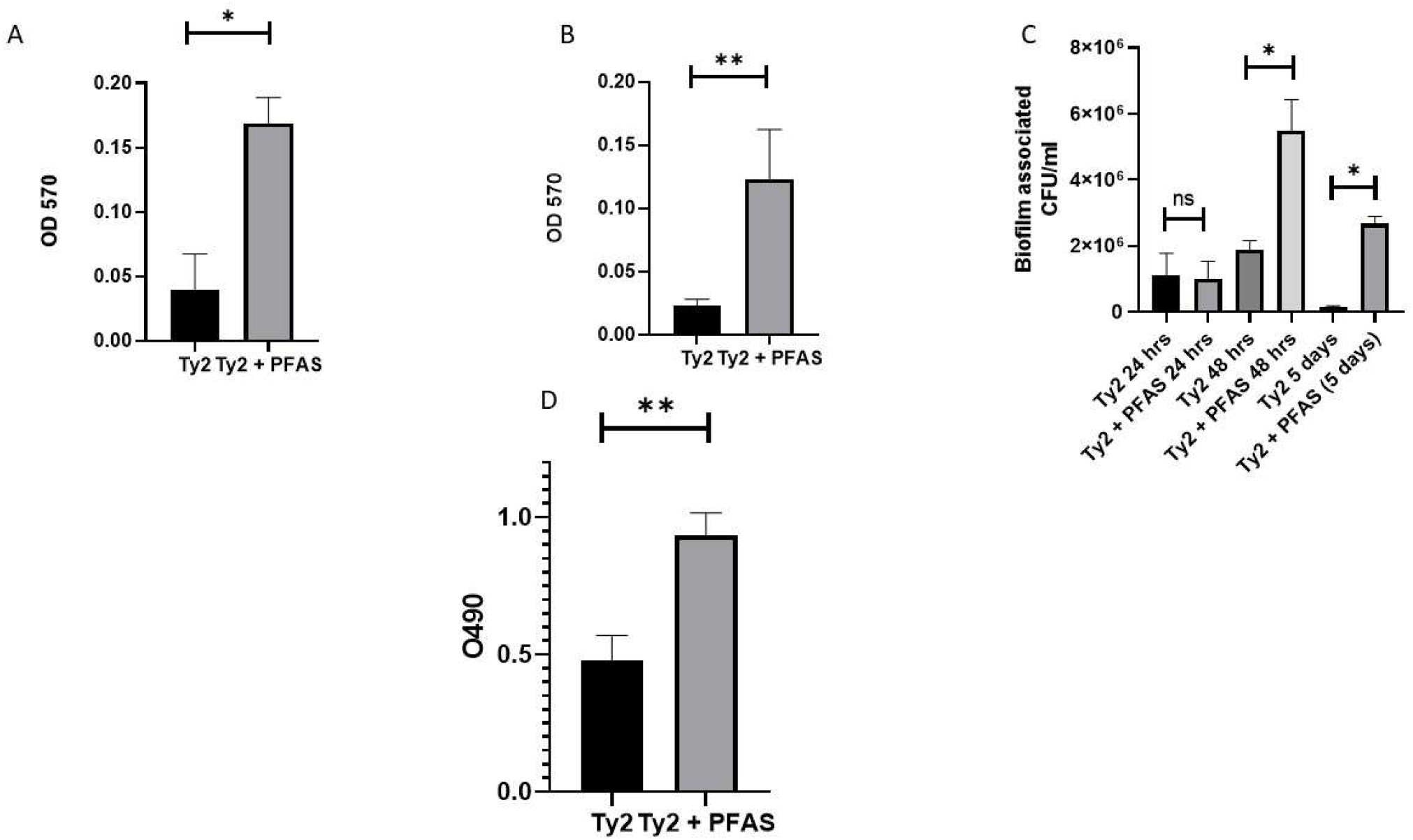
PFAS increases biofilm by Ty2: (A) About 5 × 10^9^ Ty2 cells that have been treated or not treated with PFAS were grown at 37°C in LB supplemented with streptomycin (50 µg/mL) and 2.5% ox bile for 5 days in glass tubes under stationary conditions. The biofilms obtained were then washed using PBS, stained using crystal violet dye (15 min room temperature), and the dye extracted using 50% acetic acid. (B) Biofilm formation under nutrient-limited conditions was assessed by incubating PFAS treated or untreated Ty2 cells in LB diluted with PBS (1:10) supplemented with streptomycin for 5 days on 24 wells cell plates, followed by crystal violet staining and quantification as described above. (C) Biofilm formation on biotic surfaces was determined by using sterilized lettuce leaves incubated with PFAS treated or untreated Ty2 cells in diluted LB under static conditions for 5 days. After washing, biofilms were enzymatically treated (Proteinase K and DNase I), mechanically disrupted, and viable bacteria were determined by plating serial dilutions on streptomycin-containing agar. (D) The measurement of EPS production in the presence and absence of PFAS was done using the phenol-sulphuric acid method. Data represent mean ± SD from three independent biological replicates (n = 3). Statistical significance was determined using a Mann Whitney Test, with * representing P value < 0.05 and *\*\** representing P value < 0.005.

In order to simulate environmental conditions of biofilm formation, a lettuce leaf colonization model was developed with nutrient-depleted conditions (LB at 1:10 dilutions). While no changes in the biofilm-associated bacterial numbers were observed at 24 and 48 h of cultures, biofilm-associated colony-forming units (CFU) were significantly increased when *S*. Typhi was exposed to PFAS. Moreover, biofilm associated-CFU were considerably higher even after 5 days, although total CFU were decreased (Fig. 2C). This was reflected by greater yellowing of the lettuce leaf after 5 days, indicating tissue damage (Fig. S3). In addition, EPS production was greatly increased in presence of PFAS, indicating increased biofilm formation (Fig. 2D).

### PFAS exposure reduces efflux activity and promotes an energy inefficient state in *Salmonella* Typhi

Bacterial persisters are well known to have reduced intracellular ATP owing to the inhibition of core metabolism (19). To investigate whether PFAS treatment alters cellular energetics, intracellular ATP concentrations of *S*. Typhi were measured by luminol assay. As is evident from Fig 4A, PFAS treatment significantly reduced ATP as compared with the untreated bacteria.

**Figure 3:**
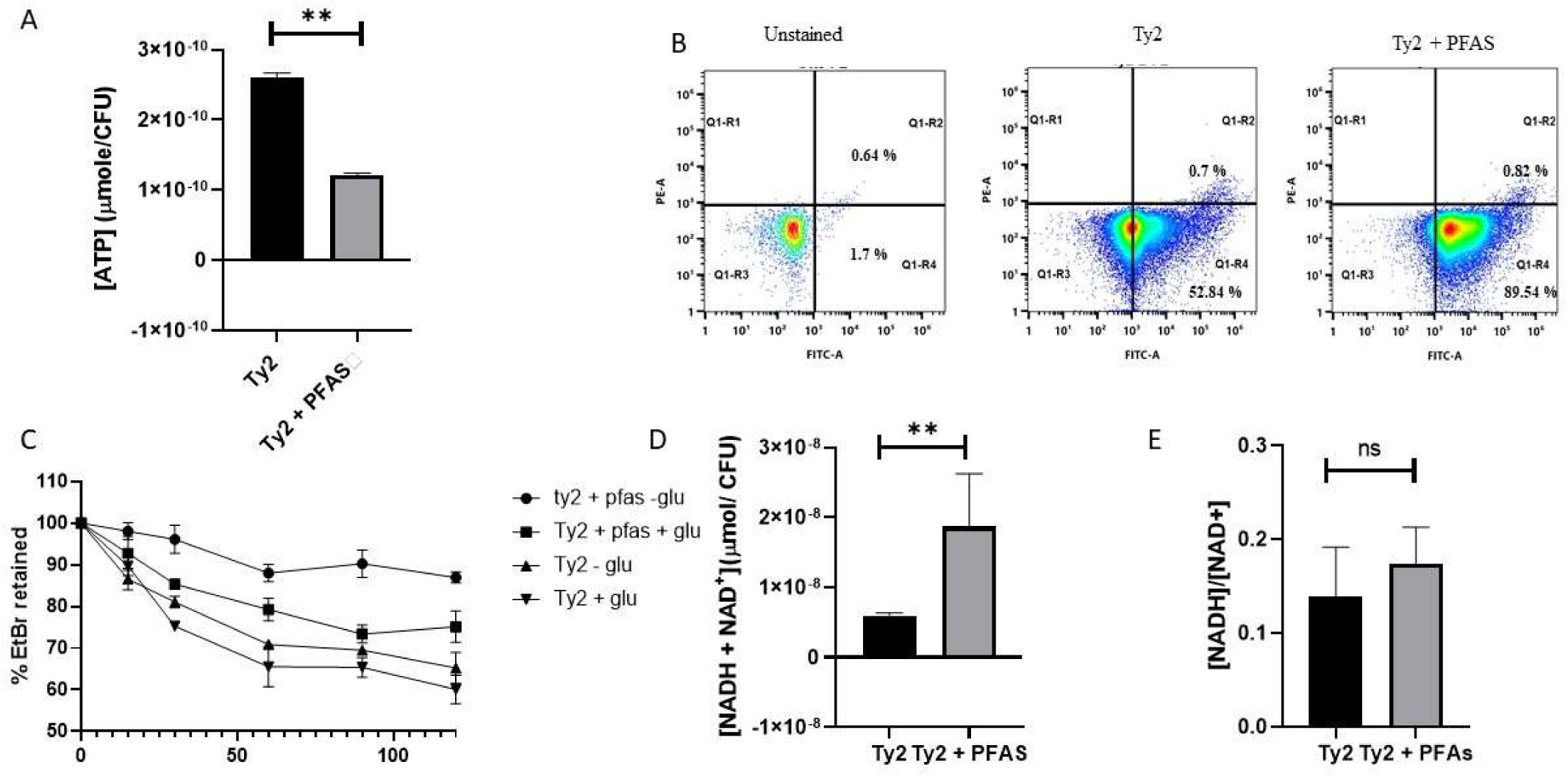
PFAS treatment increases ATP, depolarizes bacterial membrane and increase total NAD+ and NADH: (A) PFAS exposed or unexposed Ty2 cells were diluted 100-fold in PBS and placed into 96-well plates. BacTiter-Glo reagent was added, incubated for 5 min at room temperature, and luminescence was recorded to assess cellular ATP levels. (B) Membrane potential was examined by staining dilutions of the bacterial cells with DiBAC4(3) dye (1 µM, 30 min at 37°C) and subsequently analyzing them using flow cytometry. (C) The efflux activity was determined by means of ethidium bromide (EtBr) uptake and efflux assay. The cells were loaded with EtBr (1 µg/mL) and then washed, followed by resuspension in PBS, either in the presence or absence of glucose (0.4%). (D) Intracellular concentrations of NAD⁺ and NADH in PFAS-exposed and non-exposed Ty2 cells were determined through the use of a commercially available test kit, according to the kit’s guidelines. (E) The NADH / NAD⁺ ratio was determined from the above experiment (D). Data represent mean ± SD from three independent biological replicates (n = 3). Statistical significance was determined using a Mann Whitney Test, with *\*\** representing P value < 0.005.

**Figure 4:**
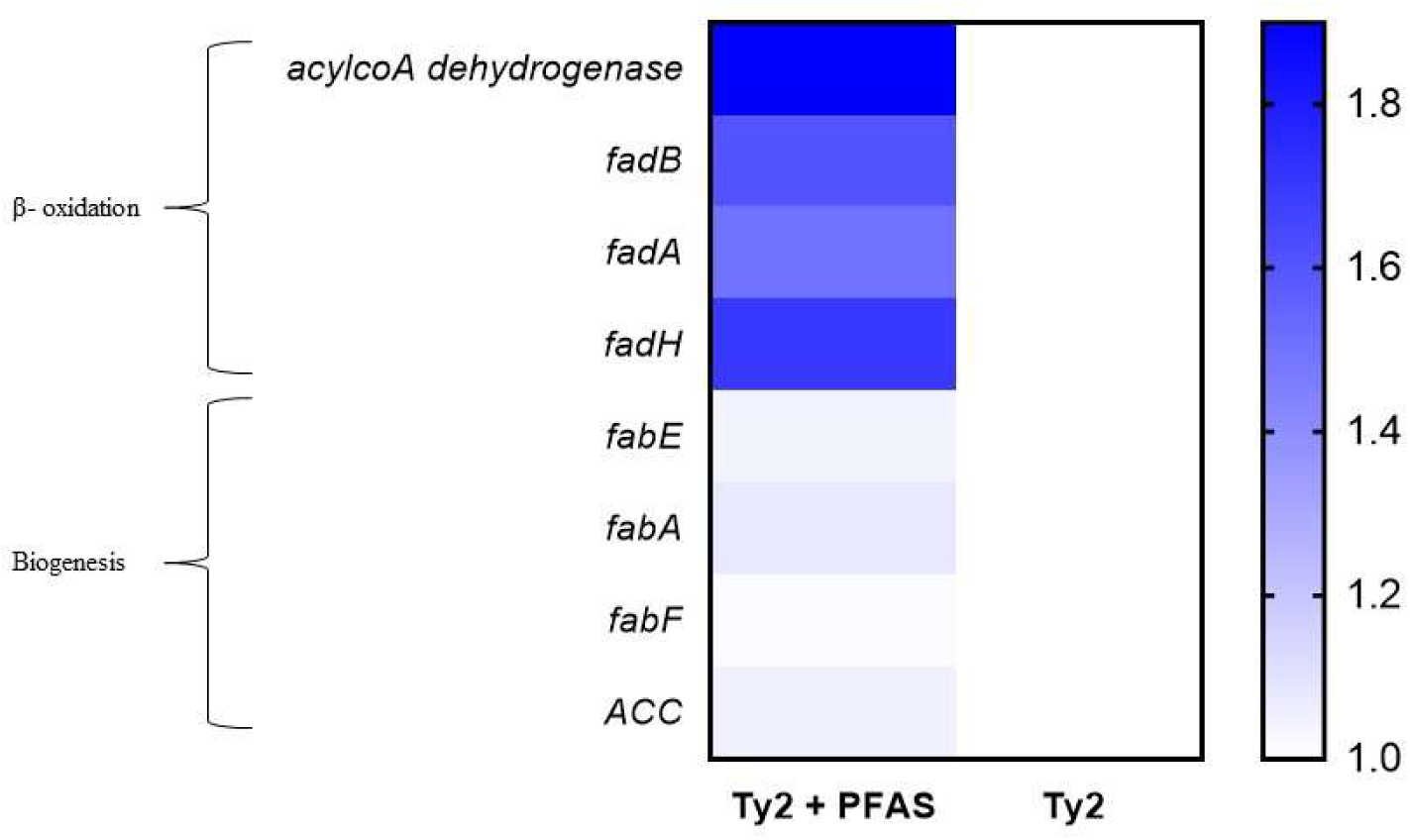
PFAS treatment enhances beta oxidation pathway without altering lipid biosynthesis pathway: Ty2 was exposed to PFAS as described in Materials and Methods. RNA was isolated from PFAS exposed or unexposed Ty2 followed by cDNA synthesis which was subsequently used for qPCR using gene specific primers. Fold change of expression of different genes involved in lipid metabolism relative to PFAS untreated control were measured using ΔΔCt method. Data represent the mean ± SD of three biological replicates (n = 3). Data represent mean ± SD from three independent biological replicates (n = 3).

Prior studies showed that PFAS treatment significantly alters bacterial membrane structures. This could result in membrane depolarisation, leading to inactivation of the membrane-bound ATP synthase. Flow cytometry analysis showed that PFAS treatment significantly increased DIBAC4(3) positive cells (52.84 % to 89.54 % positivity after PFAS), suggesting membrane depolarisation (Fig 4B). However, this was not accompanied by any significant membrane damage, leading to negligible PI positivity of the bacterial cells.

To further investigate the membrane changes, efflux pump activity of *S*. Typhi was studied after staining with EtBr. Membrane depolarisation often inhibits bacterial efflux pumps, resulting in failure of EtBr exclusion by the bacteria. As is evident from Fig 4C, PFAS treatment impaired efflux, leading to a higher percentage of EtBr+ cells, even after 120 min as compared with the PFAS-untreated controls. Interestingly, 0.4% glucose supplementation partially rescued the pump functions, further indicating that PFAS inhibited efflux pumps metabolically rather than by suppression of gene expression.

Membrane depolarisation is believed to reduce ATP synthase function by inhibiting Electron Transport Chain (ETC). To investigate this issue, total NADH+NAD^+^ of bacterial cultures was measured. Interestingly, PFAS treatment increased the total redox (NADH+NAD^+^) pool, and maintained a stable redox ratio (Fig 4D and E). This underscores a sustained or enhanced redox cycling by PFAS, while reducing ATP synthesis.

Collectively, these results suggest that PFAS disrupts the coupling of redox reactions and ATP production. This is the characteristic of an uncoupling, where energy transduction is inhibited due to the loss of the proton motive force that is required for efficient ATP production.

### PFAS induced uncoupling leads to increased lipid catabolic flux

Uncoupling of ATP synthesis and ETC is often perceived as an energy deprived state, accompanied by cellular membrane stress. Cells typically compensate this by enhancing the lipid catabolic flux to compensate the increased NAD^+^ production by ETC. To investigate the mechanisms underlying increased lipid oxidation, transcript levels of genes involved in beta oxidation and lipid biogenesis were monitored. As presented under Fig 5A, PFAS treatment significantly enhanced transcription of all the genes involved in beta oxidation pathway (*acyl coA dehydrogensase*, *fadA*, *fadB* and *fabH*), whereas major genes involved in lipid biosynthesis pathway remained unaltered (Fig 5B). These results suggest that PFAS exposure led to increased lipid catabolism within the cell, possibly in an effort to compensate for lowered ATP production to meet up the demands for continued redox reactions and to maintain membrane homeostasis. The cumulative effect was the creation of a starvation-like condition for cells, which activated the energy production pathways.

**Figure 5:**
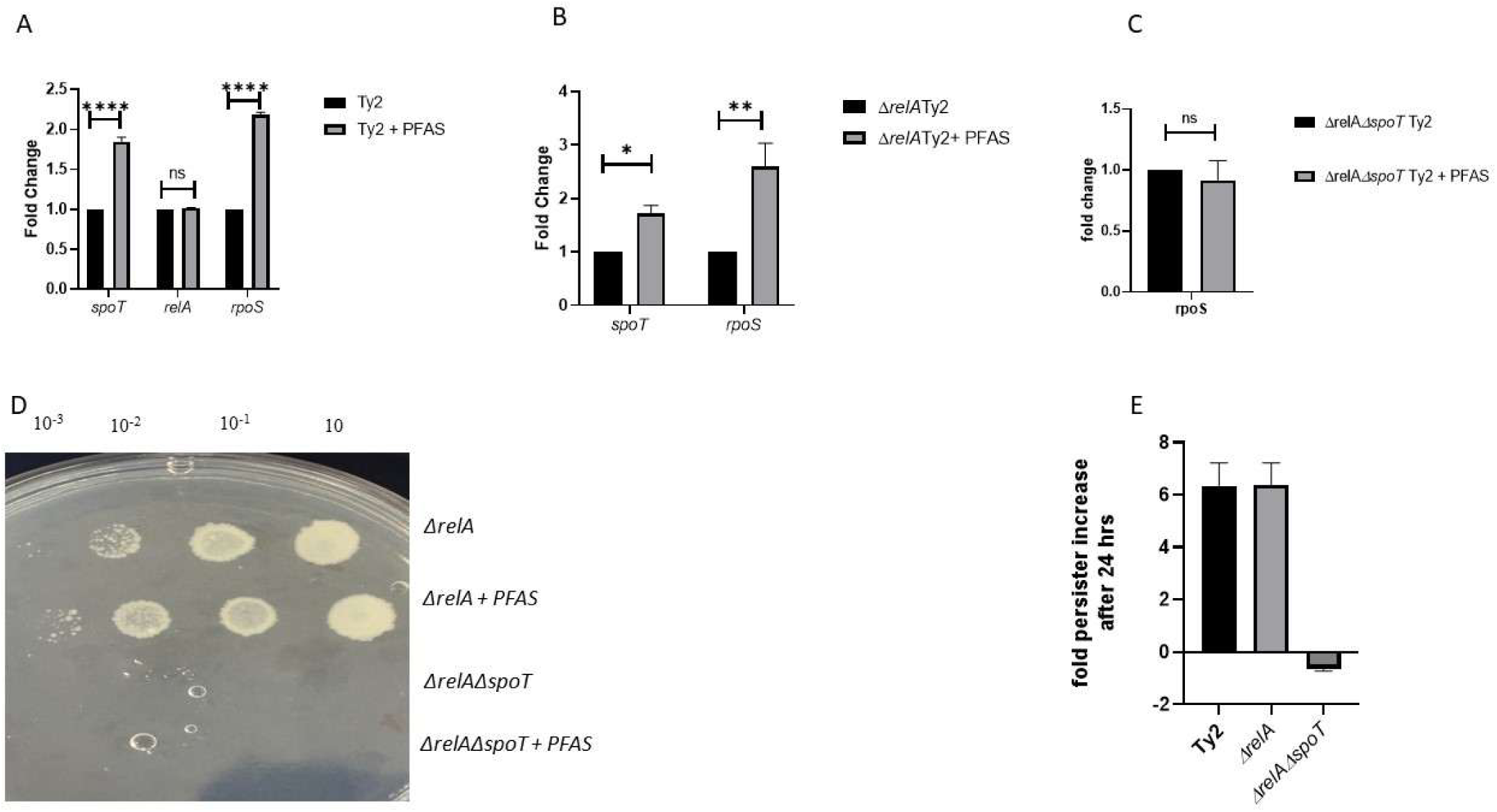
PFAS induces spot dependent stringent response: (A-C) Wild-type Ty2 (A), Δ*relA* (B), and Δ*relA*Δ*spoT* (C) strains were exposed to PFAS as described in Methods. Total RNA was isolated followed by cDNA synthesis from equal amounts of RNA. Gene expression levels were quantified by qPCR using gene-specific primers, and fold changes relative to untreated controls were calculated using the ΔΔCt method. Data represent mean ± SD from three independent biological replicates (n = 3). Statistical significance was determined using 2 way ANOVA and Wilcoxon test. **** represents < 0.0005, *** represents <0.005 and * represents < 0.05 (D) Growth phenotypes of Δ*relA* and Δ*relA*Δ*spoT* strains after PFAS exposure were determined by spot dilution assay on SMG agar plates after serial dilution. (E) Persistence following antibiotic challenge was evaluated by exposing PFAS-treated or untreated wild-type, Δ*relA*, and Δ*relA*Δ*spoT* cells (∼5 × 10⁹) to ceftriaxone (10 µg/mL) for 24 h. Surviving cells were recovered, plated on streptomycin-containing agar, and persister levels were quantified as fold change relative to untreated controls. Data represent mean ± SD from three independent biological replicates (n = 3). Statistical significance was determined using a Mann Whitney Test, with *\*\** representing P value < 0.005.

### PFAS induces a SpoT-dependent stringent response that drives RpoS activation and bacterial persistence

Energy deficient state and metabolic rewiring induced by PFAS is ideal for the activation of stringent response pathways, producing (p)ppGpp (8). To address this issue, mRNA expression of the genes regulating (p)ppGpp biosynthesis was studied by qPCR. It turned out that PFAS treatment triggered an elevation of *spoT* mRNA, while *relA* mRNA remained unchanged (Fig. 5A). Additionally, stress sigma factor *rpoS* mRNA was upregulated, pointing towards activation of a stress-related program, associated with the stringent response.

Next, stringent response independent of the *relA* gene was examined. As shown in Fig. 5B, even in the absence of *relA*, PFAS induced higher mRNA levels of both *spoT* and *rpoS*, showing that activation of the stringent response does not require RelA. In contrast, when Δ*relA* Δ*spoT* double mutant was used, PFAS failed to induce *rpoS* mRNA (Fig. 5C).

This is consistent with the observations made from the functional assay using SMG plates, where the Δ*relA* mutant displayed better growth in the presence of PFAS (Fig. 5D), suggesting an elevated production of (p)ppGpp by SpoT. PFAS treatment was found to elicit an increase in the number of persister cells resistant to ceftriaxone in the Δ*relA* mutant similar to that in the wild-type bacteria. On the other hand, there was no such elevation in persister formation in the Δ*relA*Δ*spoT* mutant upon PFAS treatment, implying SpoT’s involvement in persister cell induction (Fig 5E).

## Discussion

Environmental contaminants are increasingly acknowledged as modulators of bacterial physiology that can indirectly affect antibiotic efficacy. Per- and polyfluoroalkyl substances (PFAS) have become persistent pollutants that significantly affect microbial communities, leading to increased antimicrobial resistance (AMR) determinants and modified cellular physiology (20, 21). In this study, we illustrate that PFAS exposure reconfigures *Salmonella* Typhi into a persistence-dominant state by disrupting membrane bioenergetics, thereby offering a mechanistic framework that enhances ecological observations associating PFAS with antimicrobial resistance (AMR).

One of the key finding of this study was that there was a significant decrease in intracellular ATP levels after exposure to PFAS while preserving the amount of the total NADH + NAD^+^ pools as well as the redox ratios remaining relatively constant. This phenotype could be characterized by classical uncoupling of respiration, whereby electrons continue through ETC while the proton-motive force (PMF) is being dissipated and ATP cannot be generated efficiently (22). The collapse of electrical potential (also referred to as membrane depolarization) is supported through the significant deposition of DiBAC4(3) following PFAS exposure without increasing the amount of propidium iodide indicating that although there is a collapse in the electrical potential, the membrane has remained fairly intact. Although the function of PFAS compounds as potential agents causing membrane depolarization is known in several eukaryotes, it is for the first time we have shown that the same happens in bacteria as well (23). Membrane-targeting effects such as changes in fluidity and permeability due to exposure to PFAS have also been observed in bacteria found in the environment (14), suggesting that the disruption of membrane energetics may be a conserved phenomenon across species.

The lack of PMF has direct effects on the function of transport mechanisms in bacteria, especially those pumps that are powered by the energy of the proton motive force (22). Indeed, the decreased efficiency of these pumps in PFAS-exposed cells was confirmed by their increased retention of ethidium bromide. The ability to partially restore the pump activity under glucose addition suggests that the reduced activity of these mechanisms is due to energy deficiency, not the absence of appropriate transcription, which is consistent with previous findings demonstrating that the activity of efflux mechanisms depends strictly on energy metabolism (24).

Such apparent contradiction can be explained by the presence of the pseudo-starvation condition. Indeed, the expression of β-oxidation genes was found increased in PFAS-induced cells, indicating elevated flux through these pathways. Metabolic reprogramming leading to increased catabolism and NADH turnover is considered an adaptive response to the inefficient production of ATP in bacteria (25). In other words, electron transport continues to function in the cells but not coupling to the ATP synthesis anymore, thus supporting the uncoupling hypothesis.

One of the triggers of the stringent response is disruption of energy homeostasis. In *Salmonella* and its relatives, the protein SpoT monitors such disturbances and initiates (p)ppGpp accumulation (8). This metabolite induces stress adaptation and development of persistent forms in a manner dependent on the regulatory factor RpoS (26). Although (p)ppGpp levels were not directly measured here, the observed phenotype (reduced ATP, enhanced catabolism, increased persistence, and biofilm formation) is strongly consistent with activation of this pathway. Thus, PFAS appears to mimic nutrient limitation at the signaling level, even in nutrient rich conditions. Moreover, the induction of fatty acid catabolism due to the collapse of PMF also helps in sustaining a higher level of (p)ppGpp by activating the synthetase activity of Spot (8).

Enhancement of persister formation by membrane depolarisation is well documented in bacteria. It has been shown that several type I toxin antitoxin system in response to either DNA damage or elevated level of (p)ppGpp causes membrane depolarisation reducing intracellular ATP level and increasing the persister population (27, 28). However, in this study, for the first time we demonstrated that membrane depolarisation precedes (p)ppGpp elevation by causing an energy starved state which enhances the persister population.

Notably, our mechanistic observations are consistent with and complement recent environmental studies indicating that exposure to PFAS leads to an increase in the prevalence of AMR. Several studies indicate that PFAS exposure results in higher numbers of ARGs and MGEs, indicating HGT (29). It is important to highlight that exposure to PFAS has also been demonstrated to enhance membrane permeability and oxidative stress, factors that could contribute to plasmid transformation (30). Furthermore, conditions characterized by sublethal stress, such as reduced growth and ATP levels, promote competency and the uptake of DNA in various bacteria (31). While the present study does not test the effects of PFAS on transformation efficiency, the depolarized and low-ATP phenotype observed herein is highly conducive to HGT.

All these findings point to a dual role of PFAS on antimicrobial failure. On one hand, PFAS increases ARGs and drives gene transfer out in the environment; on the other, it pushes cells into an energy deficient state where bacteria persist longer. This combination is troubling; it connects short-term antibiotic tolerance with the slow buildup of lasting resistance. What’s striking is that the uncoupler-like action we see here doesn’t match the typical stress responses bacteria show to antibiotics. Instead, we can speculate that environmental pollutants trigger unique physiological routes to help bacteria survive.

In summary, this paper presents PFAS as a novel modulator of bacterial metabolism that functions as an uncoupler of respiration, triggers starvation stress, and enhances persistence. Through its combination of mechanistic microbiology with environmental science, our findings show the importance of accounting for non-antibiotic pollution as an active factor contributing to the emergence of antibiotic resistance and tolerance.

## Funding

The authors SD and DG would like to acknowledge the financial assistance provided by the Indian Council of Medical Research IMRG (Project ID: IIRP-2023-0000225) and SC would like to acknowledge UGC for PhD fellowship.

## Conflict of interest

We declare that there is no conflict of interest regarding the publication of this article.

## Supplementary Figure Legends

**Figure S1:**
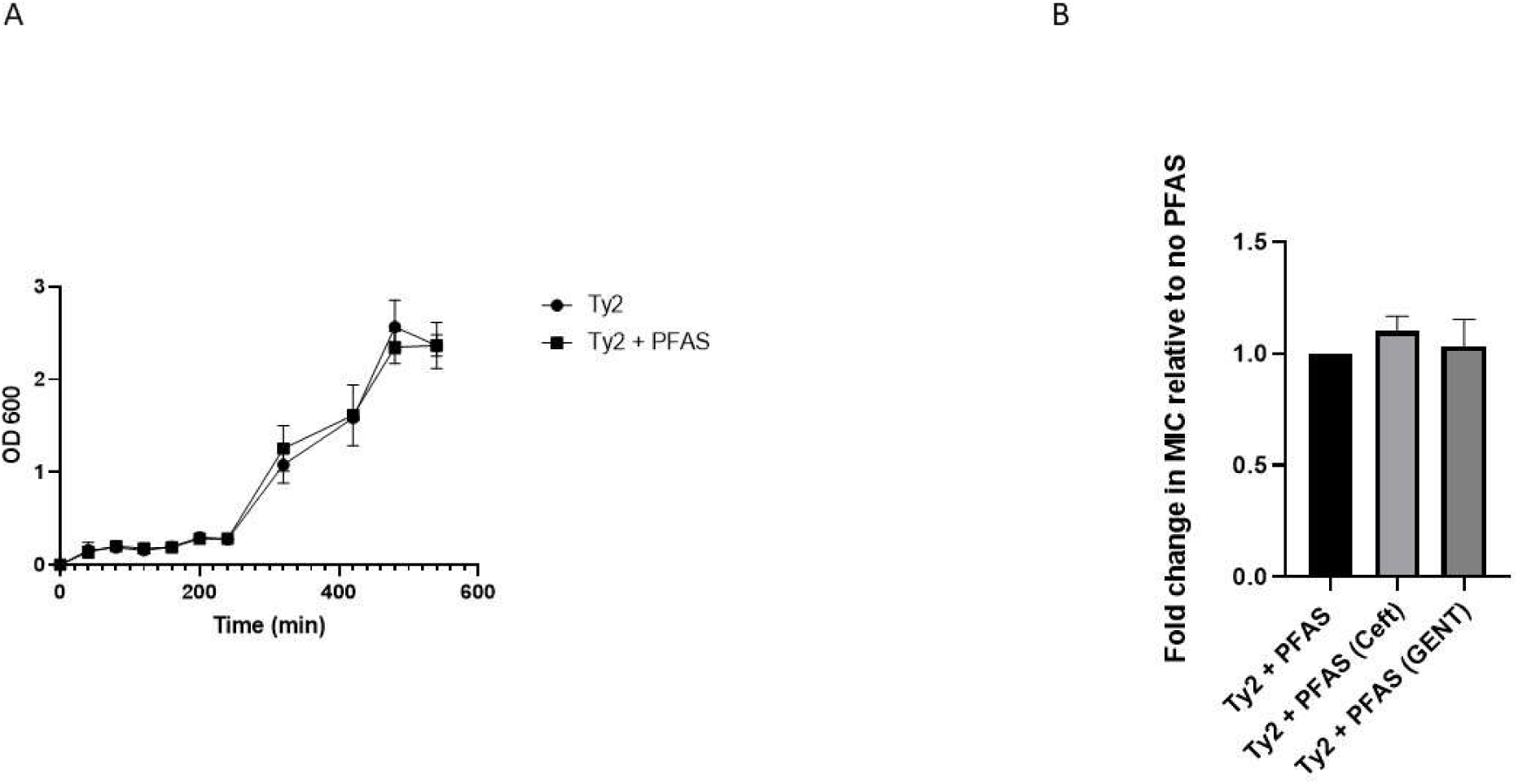
PFAS neither inhibit growth nor MIC of Ty2: (A) Overnight grown culture of PFAS treated or untreated Ty2 strain were diluted 1:100 with fresh LB media and incubated at 37C under shaking condition. Cultures were taken out at definite time intervals and OD was measured at 600 nm.(B) Overnight grown culture of PFAS treated or untreated Ty2 strain were diluted 1:100 with fresh LB media with or without increasing concentrations of antibiotics and incubated at 37C under shaking condition. Fold change of MIC for respective antibiotics relative to no PFAS control were calculated after 16 hrs of incubation. Data represent the mean ± SD of three biological replicates (n = 3). Statistical significance was determined using a two-tailed Student’s t-test. *P* < 0.005 was considered statistically significant.

**Figure S2:**
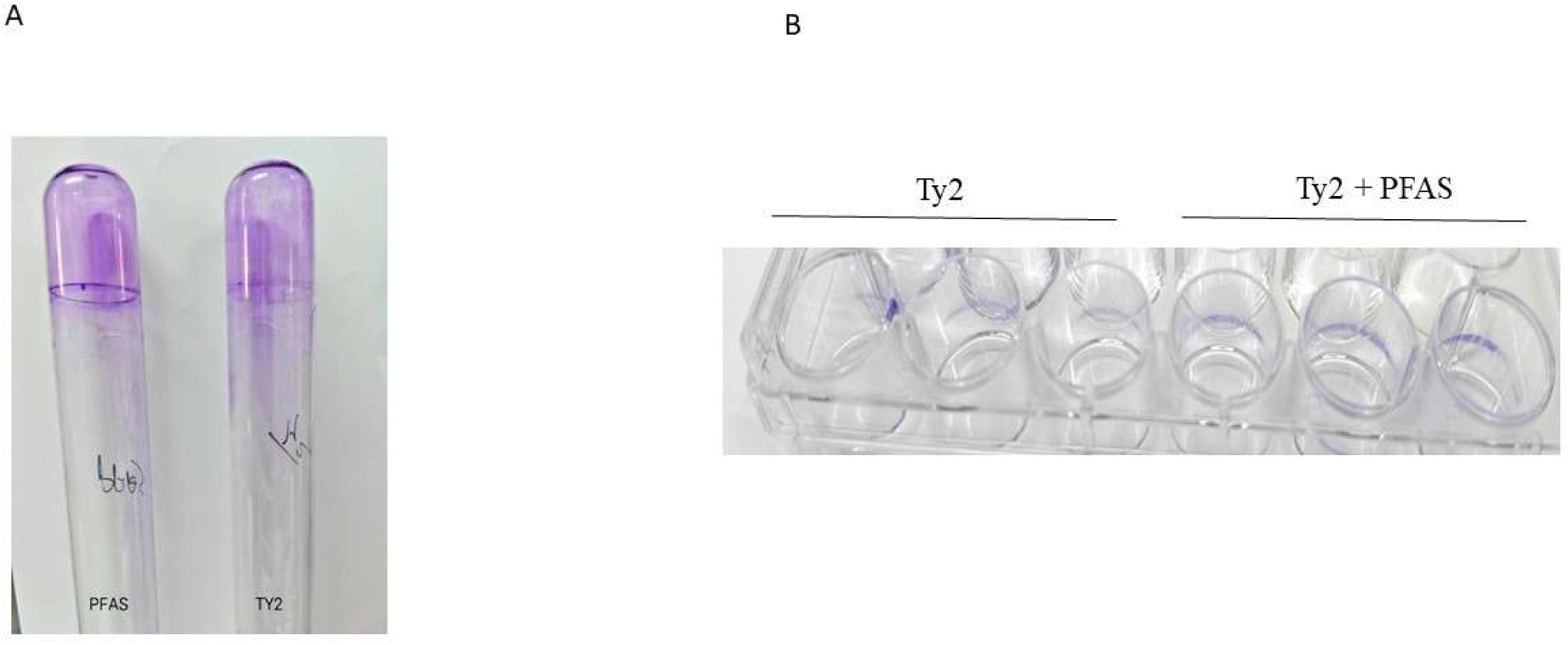
PFAS increases biofilm by Ty2. (A) Represented image showing crystal violet staining of biofilm on the test tube wall in nutrient rich LB media containing 5% ox bile of PFAS untreated and treated Ty2. (B) Represented image showing crystal violet staining of biofilm on the wall of 24 well plate in nutrient poor media of PFAS untreated and treated Ty2.

**Figure S3:**
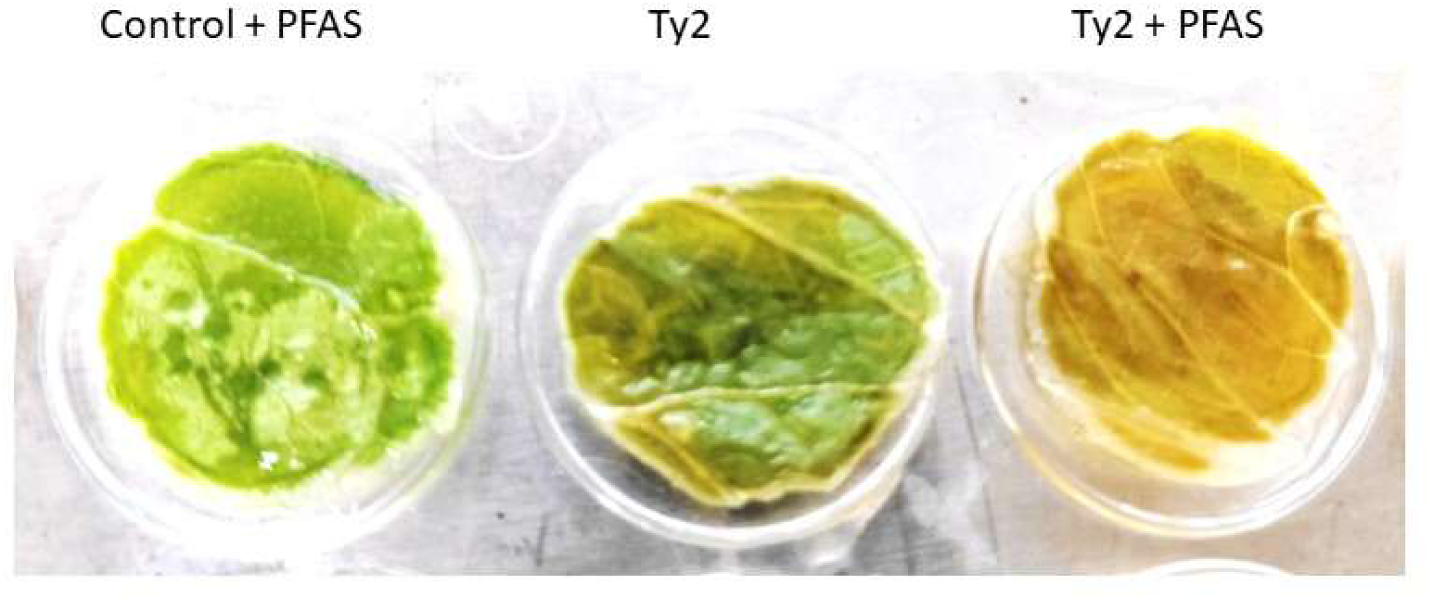
PFAS increases bacterial localization on lettuce leaf: Representative image showing chlorosis of lettuce leaf by Ty2 in the presence of PFAS. More yellow coloration indicated more chlorosis and hence more bacterial deposition.

## References

1. Crump, J. A., & Mintz, E. D. (2010). Global trends in typhoid and paratyphoid fever. Clinical infectious diseases, 50(2), 241–246. doi: 10.1086/649541.

2. Dougan, G., & Baker, S. (2014). Salmonella enterica serovar Typhi and the pathogenesis of typhoid fever. Annual review of microbiology, 68(1), 317–336. doi: 10.1146/annurev-micro-091313-103739.

3. Galán, J. E. (2001). Salmonella interactions with host cells: type III secretion at work. Annual review of cell and developmental biology, 17(1), 53–86. doi: 10.1146/annurev.cellbio.17.1.53.

4. Ibarra, J. A., & Steele-Mortimer, O. (2009). Salmonella–the ultimate insider. Salmonella virulence factors that modulate intracellular survival. Cellular microbiology, 11(11), 1579–1586. doi: 10.1111/j.1462-5822.2009.01368.x.

5. Balaban, N. Q., Helaine, S., Lewis, K., Ackermann, M., Aldridge, B., Andersson, D. I., … & Zinkernagel, A. (2019). Definitions and guidelines for research on antibiotic persistence. Nature Reviews Microbiology, 17(7), 441–448. DOI: 10.1038/s41579-019-0196-3

6. Potrykus, K., & Cashel, M. (2008). (p) ppGpp: still magical?. Annu. Rev. Microbiol., 62(1), 35–51. DOI: 10.1146/annurev.micro.62.081307.162903

7. Dalebroux, Z. D., & Swanson, M. S. (2012). ppGpp: magic beyond RNA polymerase. Nature Reviews Microbiology, 10(3), 203–212. DOI: 10.1038/nrmicro2720

8. Battesti, A., & Bouveret, E. (2006). Acyl carrier protein/SpoT interaction, the switch linking SpoT-dependent stress response to fatty acid metabolism. Molecular microbiology, 62(4), 1048–1063. DOI: 10.1111/j.1365-2958.2006.05442.x

9. Salzer, A., & Wolz, C. (2023). Role of (p) ppGpp in antibiotic resistance, tolerance, persistence and survival in Firmicutes. Microlife, 4, uqad009. DOI: 10.1093/femsml/uqad009

10. Kurata, T., & Takada, H. (2026). Mechanistic and evolutionary perspective of RelA/SpoT homologus from and beyond stringent-response signalling. Bioscience, Biotechnology, and Biochemistry, zbag003. DOI: 10.1093/bbb/zbag003

11. Parsons, J. B., & Rock, C. O. (2013). Bacterial lipids: metabolism and membrane homeostasis. Progress in lipid research, 52(3), 249–276. DOI: 10.1016/j.plipres.2013.02.002

12. Campbell, J. W., & Cronan Jr, J. E. (2001). Bacterial fatty acid biosynthesis: targets for antibacterial drug discovery. Annual Reviews in Microbiology, 55(1), 305–332. DOI: 10.1146/annurev.micro.55.1.305

13. Abunada, Z., Alazaiza, M. Y., & Bashir, M. J. (2020). An overview of per-and polyfluoroalkyl substances (PFAS) in the environment: Source, fate, risk and regulations. Water, 12(12), 3590. 10.3390/w12123590

14. Panella, M., Rabadi, A., Ceja-Vega, J., Said, J., Andersen, E., Mitchell, J., … & Lee, S. (2025). Membrane-modifying effects of perfluoroalkyl substances in model bacterial membranes. ACS omega, 10(35), 39884–39897. doi: 10.1021/acsomega.5c04177

15. Proia, L., Lupini, G., Osorio, V., Pérez, S., Barceló, D., Schwartz, T., … & Sabater, S. (2013). Response of biofilm bacterial communities to antibiotic pollutants in a Mediterranean river. Chemosphere, 92(9), 1126–1135. DOI: 10.1016/j.chemosphere.2013.01.063

16. Harrison, J. J., Ceri, H., & Turner, R. J. (2007). Multimetal resistance and tolerance in microbial biofilms. Nature Reviews Microbiology, 5(12), 928–938. DOI: 10.1038/nrmicro1774

17. Dasgupta, S., Das, S., Biswas, A., Bhadra, R. K., & Das, S. (2019). Small alarmones (p) ppGpp regulate virulence associated traits and pathogenesis of Salmonella enterica serovar Typhi. Cellular microbiology, 21(8), e13034. DOI: 10.1111/cmi.13034

18. Ganguli, D., Chakraborty, S., Chakraborty, S., Pal, A., Gope, A., & Das, S. (2022). Macrophage cell lines and murine infection by Salmonella enterica serovar typhi L-form bacteria. Infection and Immunity, 90(6), e00119–22. DOI: 10.1128/iai.00119-22

19. Shan, Y., Brown Gandt, A., Rowe, S. E., Deisinger, J. P., Conlon, B. P., & Lewis, K. (2017). ATP-dependent persister formation in Escherichia coli. MBio, 8(1), 10–1128. DOI: 10.1128/mBio.02267-16

20. Sun, H., Hu, J., Wang, B., Wang, G., Liu, Y., Yang, X., … & Zhang, S. (2025). Short-chain PFAS in coastal sediments: PFBS-driven antimicrobial resistance and pathogen risks. Water Research, 124579. DOI: 10.1016/j.watres.2025.124579

21. Zhang, T., Xu, Z., & Fang, H. H. P. (2022). PFAS exposure reshapes microbial communities and resistome profiles. Water Research, 214, 118189. 10.1016/j.watres.2022.118189

22. Klingenberg, M. (1999). Uncoupling protein—A useful energy dissepator. Journal of bioenergetics and biomembranes, 31(5), 419–430. DOI: 10.1023/a:1005440221914

23. Kam, Y., Winer, L., & Romero, N. (2025). Chain length-dependent mitochondrial toxicity of perfluoroalkyl carboxylic acids: insights from Mito Tox Index evaluation. Frontiers in Toxicology, 7, 1582891. DOI: 10.3389/ftox.2025.1582891

24. Whittle, E. E., Orababa, O., Osgerby, A., Siasat, P., Element, S. J., Blair, J. M., & Overton, T. W. (2024). Efflux pumps mediate changes to fundamental bacterial physiology via membrane potential. MBio, 15(10), e02370–24. DOI: 10.1128/mbio.02370-24

25. Black, P. A., Warren, R. M., Louw, G. E., van Helden, P. D., Victor, T. C., & Kana, B. D. (2014). Energy metabolism and drug efflux in Mycobacterium tuberculosis. Antimicrobial agents and chemotherapy, 58(5), 2491–2503. DOI: 10.1128/AAC.02293-13

26. Pacios, O., Blasco, L., Bleriot, I., Fernandez-Garcia, L., Ambroa, A., López, M., … & Tomás, M. (2020). (p) ppGpp and its role in bacterial persistence: new challenges. Antimicrobial agents and chemotherapy, 64(10), 10–1128. DOI: 10.1128/AAC.01283-20

27. Verstraeten, N., Knapen, W. J., Kint, C. I., Liebens, V., Van den Bergh, B., Dewachter, L., … & Michiels, J. (2015). Obg and membrane depolarization are part of a microbial bet-hedging strategy that leads to antibiotic tolerance. Molecular cell, 59(1), 9–21. DOI: 10.1016/j.molcel.2015.05.011

28. Berghoff, B. A., & Wagner, E. G. H. (2019). Persister formation driven by TisB-dependent membrane depolarization. In Persister cells and infectious disease (pp. 77–97). Cham: Springer International Publishing. DOI:10.1007/978-3-030-25241-0_5

29. Gang, D., Li, Z., Yu, H., Hu, C., & Qu, J. (2025). PFAS Stress on Functional Expression of Periphyton Communities and Trade-off Strategies for Horizontal/Vertical Transfer of Resistance Genes. Environmental Science & Technology, 59(24), 12255–12267. DOI: 10.1021/acs.est.5c02692

30. Li, Y., Zhang, Y., Liu, X., Zhou, X., Ye, T., Fu, Q., … & Wang, D. (2025). Per-and polyfluoroalkyl substances exacerbate the prevalence of plasmid-borne antibiotic resistance genes by enhancing natural transformation, in vivo stability, and expression in bacteria. Water Research, 272, 122972. DOI: 10.1016/j.watres.2024.122972

31. da Cunha, E. T., Pedrolo, A. M., & Arisi, A. C. M. (2023). Effects of sublethal stress application on the survival of bacterial inoculants: a systematic review. Archives of microbiology, 205(5), 190. DOI: 10.1007/s00203-023-03542-8

